# N-Power AI: A Specialized Agent Framework for Automated Sample Size and Power Analysis in Clinical Trial Design

**DOI:** 10.1101/2025.02.06.636776

**Authors:** Peifeng Ruan, Ismael Villanueva-Miranda, Jialiang Liu, Donghan M. Yang, Qinbo Zhou, Guanghua Xiao, Yang Xie

## Abstract

**Background:** Sample size and power analysis are essential in biomedical research and investigations, particularly in clinical trial design, as they ensure sufficient statistical power to detect meaningful effects. However, the complexity of these calculations often requires specialized statistical expertise, making the process inconvenient and limiting accessibility for researchers during early-stage study planning.

**Methods:** We developed N-Power AI, an agentic framework leveraging large language models (LLMs) to perform sample size and power calculations across diverse study designs. The framework consists of three specialized agents: the Function Agent, a perception and reasoning module that identifies appropriate statistical tests and corresponding R functions; the Calculation Agent, an action module that extracts parameters and executes precise computations; and the Reporting Agent, a presentation module that generates comprehensive, downloadable reports. N-Power AI and advanced LLMs (e.g., GPT-o1, Claude 3.5, Gemini 1.5 Pro) were evaluated against ground truths from statistical software (R) across six common clinical trial scenarios.

**Results:** Direct LLM outputs showed significant deviations from ground-truth values, particularly in complex scenarios like the Chi-Square Test and Cox Proportional Hazards Model. In contrast, N-Power AI achieved 100% agreement with ground truths across all scenarios. This accuracy is attributed to the Function Agent’s correct selection of statistical methods, the Calculation Agent’s accurate computations, and the Reporting Agent’s ability to produce clear and comprehensive summaries.

**Conclusion:** N-Power AI automates sample size and power analysis, offering an effective, efficient, and accessible solution for early-stage study planning. While human expertise remains crucial for high-level statistical planning, N-Power AI enhances accessibility and efficiency, streamlining the analysis process to generate reliable and reproducible results for a wide range of research scenarios.

## Introduction

An effective statistical analysis plan is the biomedical research and investigations, particularly in clinical trial design, ensuring valid, reliable, and replicable results. Questions regarding the quantitative properties of clinical trial designs, especially the appropriate power and optimal sample size, are among the most frequently asked by clinicians to statisticians^1^. The significance of such power analysis, including calculating the necessary sample size for a specified power and calculating the power when given a specific sample size, spans various study designs, affecting the credibility and reproducibility of results in clinical research^2^. Specifying these quantitative properties of a clinical trial requires both conceptual and computational effort, which involves considering study design, endpoints, assumptions, and statistical methods^1^. The complexity of these calculations often demands specialized statistical expertise, which may not be readily available to all researchers. This challenge is intensified by the growing need for thorough statistical planning, driven by the increasing complexity of clinical trials and heightened regulatory oversight. While powerful tools for sample size and power calculations have been developed, such as R^3^, SAS^4^ or specialized tools like G^*^ Power^5^, they often require a steep learning curve and a deep understanding of statistical theory. Consequently, there is an urgent demand for more accessible and efficient solutions that can facilitate statistical planning while ensuring accuracy and reliability.

Recent advancements in large language models (LLMs) have created new opportunities for automating complex tasks in fields like medical research^6^. LLMs excel in natural language processing, data analysis, and code generation, making them potential tools for streamlining statistical workflows^7^. However, the ability of LLMs to perform mathematical computations and follow statistical principles, which are crucial for accurate sample size and power calculations, is still unclear. While there have been recent investigations into the use of LLMs for matching patients to clinical trials^8^, evaluating medical adherence to guidelines in trials^9^, and autonomously answering clinical questions^10^, we still lack evidence on whether and how LLMs might be employed for sample size and power calculations.

In this work, we evaluated the ability of various cutting-edge LLMs (Llama 3.1, GPT-o1, GPT-4o, GPT-4o mini, Claude 3.5, Gemini 1.5 Pro, and GROK 2) to conduct power analyses. We compared their outputs to ground truths from statistical software (R) across six common clinical trial scenarios, assessing their accuracy in sample size and power calculations. Deviances from the ground truth were observed in sample size and power calculations when directly utilizing various LLMs, especially in more complex scenarios. We then introduce N-Power AI, an automated tool that leverages a specialized framework built on LLMs (Figure 1) to assist researchers in conducting power analyses for various clinical trial designs. N-Power AI employs a three-agent framework: a Function Agent as the perception and reasoning module to identify appropriate statistical tests and corresponding R functions, a Calculation Agent as the action module to extract parameters and perform precise calculations, and a Reporting Agent as the presentation module to generate comprehensive, downloadable reports, ensuring that the tool maintains the accuracy and reliability of traditional statistical software while offering the accessibility and efficiency of LLM-driven analysis automation.

**Figure 1.**
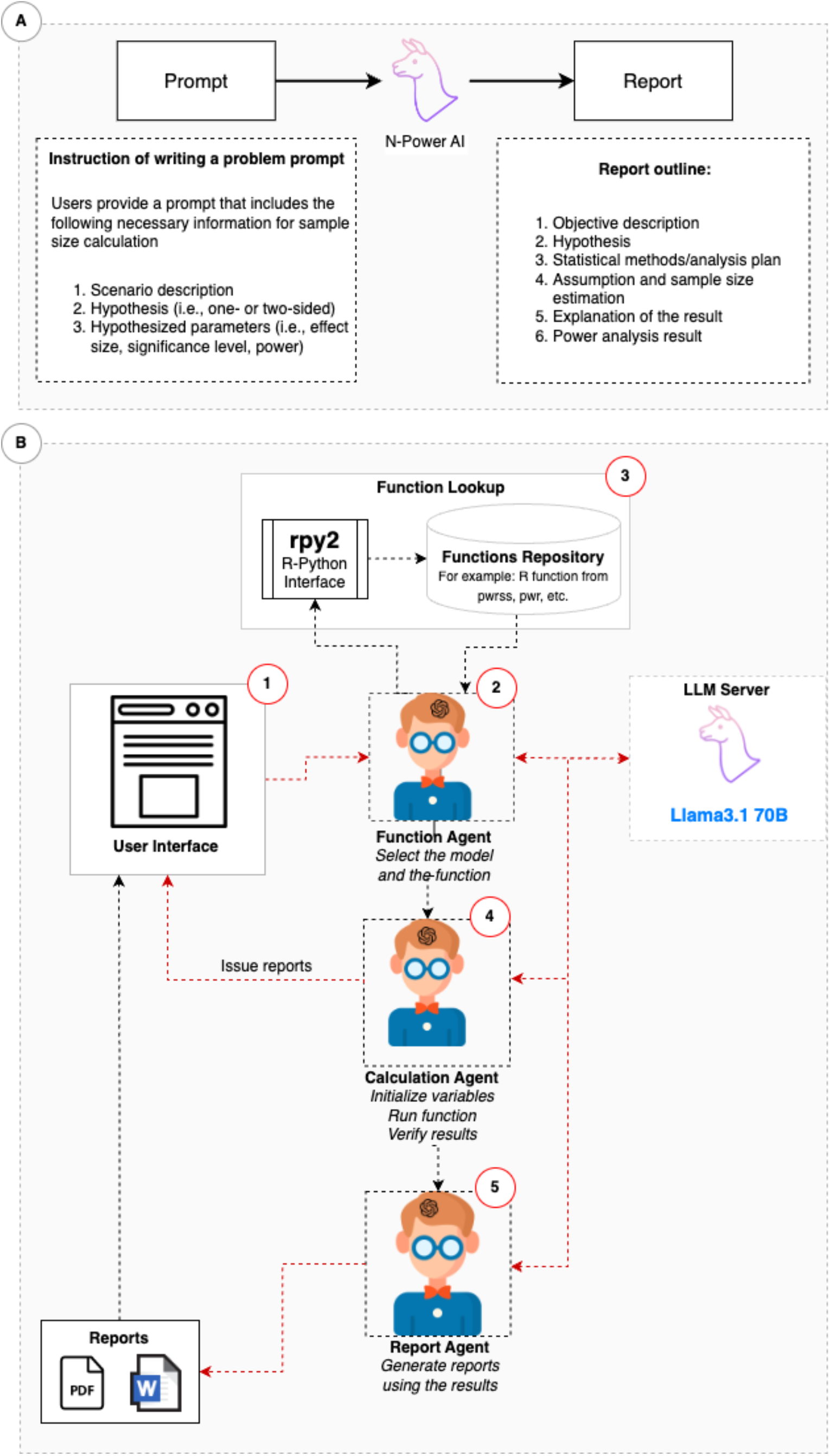
Workflow of N-Power AI. A). The proposed N-Power AI is designed to process natural language inputs containing essential details, such as scenario descriptions, hypotheses, and parameters, delivering accurate sample size or power calculations to users. B). The N-Power AI framework consists of three specialized LLM-powered agents, each dedicated to specific tasks to ensure accurate and efficient sample size calculation and power estimation. These agents collaborate by exchanging information to complete the sample size or power calculation based on user input. Specifically, the framework includes: (1) Function Agent: this agent identifies appropriate statistical tests and corresponding R functions based on the input scenario. (2) Calculation Agent: this agent extracts relevant parameters and calculates sample size and power estimates using the selected R function. (3) Reporting Agent: this agent generates a downloadable report summarizing the results, including key calculations and methodological details.

## Methods

### THE THREE-AGENT FRAMEWORK

The proposed N-Power AI is designed to process natural language inputs containing essential details, such as scenario descriptions, hypotheses, and parameters, and deliver accurate sample size or power calculations to users (Figure 1a). N-Power AI framework consists of three specialized LLM-based agents, each dedicated to specific tasks to ensure accurate and efficient sample size calculation and power estimation, as illustrated in Figure 1b. AI agents are defined as a class of interactive systems that can perceive inputs, such as natural language, visual stimuli, and other environmentally grounded data, and can produce meaningful embodied actions^11^. Specifically, LLM-based agents integrate LLMs with specialized components such as planning modules and memory mechanisms to execute complex, multi-step tasks. These agents can dynamically adapt to new information, retain context across interactions, and make informed, context-aware decisions, enabling them to handle intricate workflows with precision and reliability^12^. When building applications with LLMs, agents are the better option when flexibility and model-driven decision-making are required^13^. This is especially critical in tasks like conducting power analyses, where LLM-based agents are essential to address the limitations of direct LLM calls in handling complex mathematical computations and domain-specific requirements. These agents collaborate, exchanging information to complete the sample size or power calculation with user input. Specifically, the framework includes:

#### Function Agent

Identifies appropriate statistical tests and the corresponding R functions based on the user’s input scenario.

#### Calculation Agent

Extracts relevant parameters and calculates sample sizes and power estimates using the selected R function.

#### Reporting Agent

Generates a downloadable report summarizing the results, including key calculations and methodological details.

The framework was developed in Python, leveraging the rpy2 library^14^ to integrate Python’s robust ecosystem and the availability of high-level APIs for LLMs with R’s comprehensive statistical package. The LLM server was built using the Ollama library^15^ with the Llama3.1 model^16^, while OpenAI’s API^17^ facilitated communication among the agents.

### THE FUNCTION AGENT

The Function agent serves as a perception and reasoning module within the N-Power AI framework, responsible for receiving and processing natural language inputs, as well as conducting logical reasoning and decision-making. When a user submits a study description through the web interface (Figure 1b, step 1), the function agent connects with the LLM server and selects an appropriate statistical method for the study (Figure 1b, step 2). Next, the Function agent chooses an R function from a predefined repository, such as the pwrss package^18^ (Figure 1b, steps 3). This ensures that the selected statistical method is accurately mapped to the appropriate computational tool. Finally, the Function Agent sends the chosen R function to the Calculation Agent, guaranteeing that the user’s study is aligned with the correct statistical test and ready for further analysis by the Calculation Agent.

### THE CALCULATION AGENT

The calculation agent serves as an action module within the N-Power AI framework, responsible for executing the R function chosen by the function agent (Figure 1b, step 4) using the function calling approach. Function calling refers to the capability of integrating LLMs with external tools, like Python and R, ensuring effective utilization and seamless interaction with external APIs^19^. To achieve this, the Calculation Agent connects to the LLM server to extract necessary parameters from the users’ input, including effect size, significance level, power, alternatives, etc. Afterward, the selected R function is executed by the Calculation Agent with the parameters. Once the execution is complete, the calculation agent reviews the calculations to verify the outputs by checking whether the input and output of each step align with the function’s requirements. If any issues, such as parameters falling outside a reasonable range (e.g., probabilities not within 0 and 1) or unexpected data types (e.g., characters used for numerical inputs) are detected, the agent provides detailed issue reports, describes the issue, explains its potential impact on the analysis, and provides actionable suggestions for resolving the problem.

### THE REPORTING AGENT

The reporting agent acts as a presentation module within the N-Power AI framework and is responsible for compiling the findings into a comprehensive and user-friendly report (Figure 1b, step 5). The report is designed to provide a clear and structured summary of the analysis, including the study objectives, hypotheses, selected statistical models, underlying assumptions, and detailed analysis results (Figure 2, S1 – S5). The final report is available for download in both PDF and Word formats, allowing users to easily share, print, or integrate the results into their research documentation.

**Figure 2.**
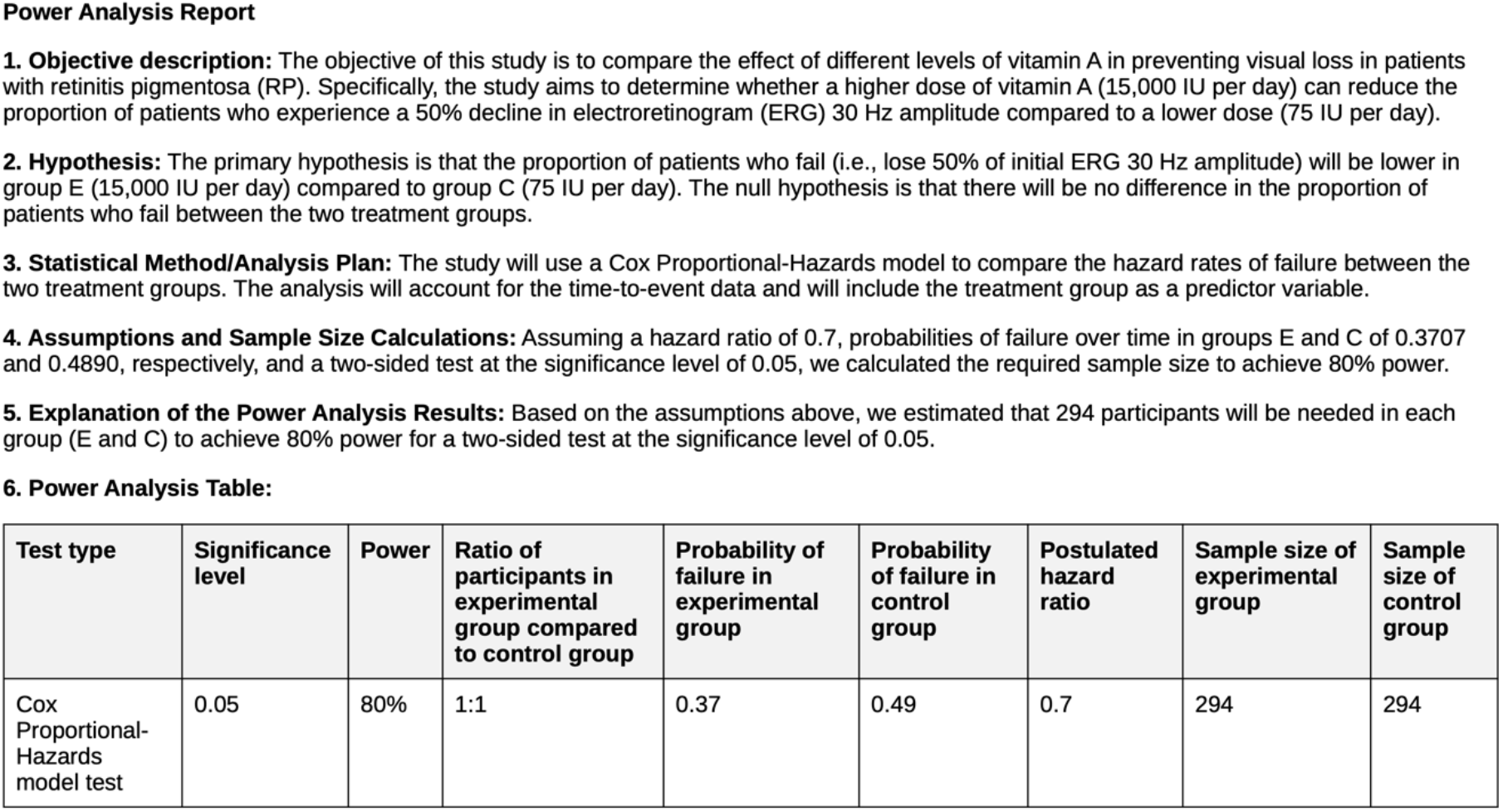
Example report generated by N-Power AI for the Cox Proportional-Hazards model scenario

## Results

### EXPERIMENTS SETTINGS

To evaluate the performance of direct LLM calls and the proposed N-Power AI framework, we selected six common scenarios in clinical trial design: (1) assessing whether the mean of a sample differs from a known or target value (one-sample t-test), (2) comparing the means of two independent groups (two-sample t-test), (3) evaluating differences in means between paired samples (paired t-test), (4) determining whether at least one group differs significantly across three or more independent groups (one-way ANOVA), (5) examining associations between categorical variables (chi-square test), and (6) analyzing the relationship between predictor variables and time-to-event data (Cox proportional hazards model). Each scenario was generated by revising cases from established sources, such as textbook examples^20,21,^ published research^22^, and online course materials^23^. These scenario descriptions were used as input for various advanced LLMs and the N-Power AI framework. The detailed scenario descriptions are listed in Table S1.

For each scenario, ground-truth sample sizes and power estimates were calculated using R 4.2.2. We then employed advanced LLMs, including GPT-o1^24^, GPT-4o^25^, GPT-4o-mini^26^, Llama 3.1^16^, Claude 3.5^27^, Gemini 1.5 Pro^28^, and GROK 2^29^ to calculate sample sizes and estimate statistical power for each scenario. The N-Power AI framework was implemented under various LLM configurations, assigning the agents’ tasks to different LLMs, including Llama 3.1 70b, Llama 3.1 70b-instruct-8k, GPT-4o-mini, and GPT-4o. For every scenario and LLM setting, we ran the N-Power AI ten times to compute sample sizes or power estimates. We compared the results to the ground truths generated by R to evaluate the performance of the direct LLMs calls and the proposed N-Power AI across various clinical trial design contexts.

### NECESSITY OF SPECIALIZED AGENT FRAMEWORK

Is it possible to calculate sample sizes and powers by directly asking an LLM model? To answer this question, we tested the performance of direct LLM calls (without using our specialized agent framework) for calculating sample sizes and power estimates. Using the same six scenarios from Table S1 as input, we asked various LLM models^24-29^ to compute sample sizes or power estimates.

Table 1 shows sample size calculations across the six scenarios, demonstrating the performance of various large language models (LLMs) compared with ground truth values derived from R. Cutting edge LLMs, including GPT-o1^24^, GPT-4o^25^, GPT-4o-mini^26^, Llama 3.1^16^, Claude 3.5^27^, Gemini 1.5 Pro^28^, and GROK 2^29^, were evaluated on their ability to accurately estimate sample sizes. All seven LLMs’ results showed deviations from the ground truth sample sizes derived from R crossing all six scenarios, especially for more complex scenarios when statistical models such as the Chi-Square Test and Cox Proportional Hazards Model are needed.

**Table 1.**
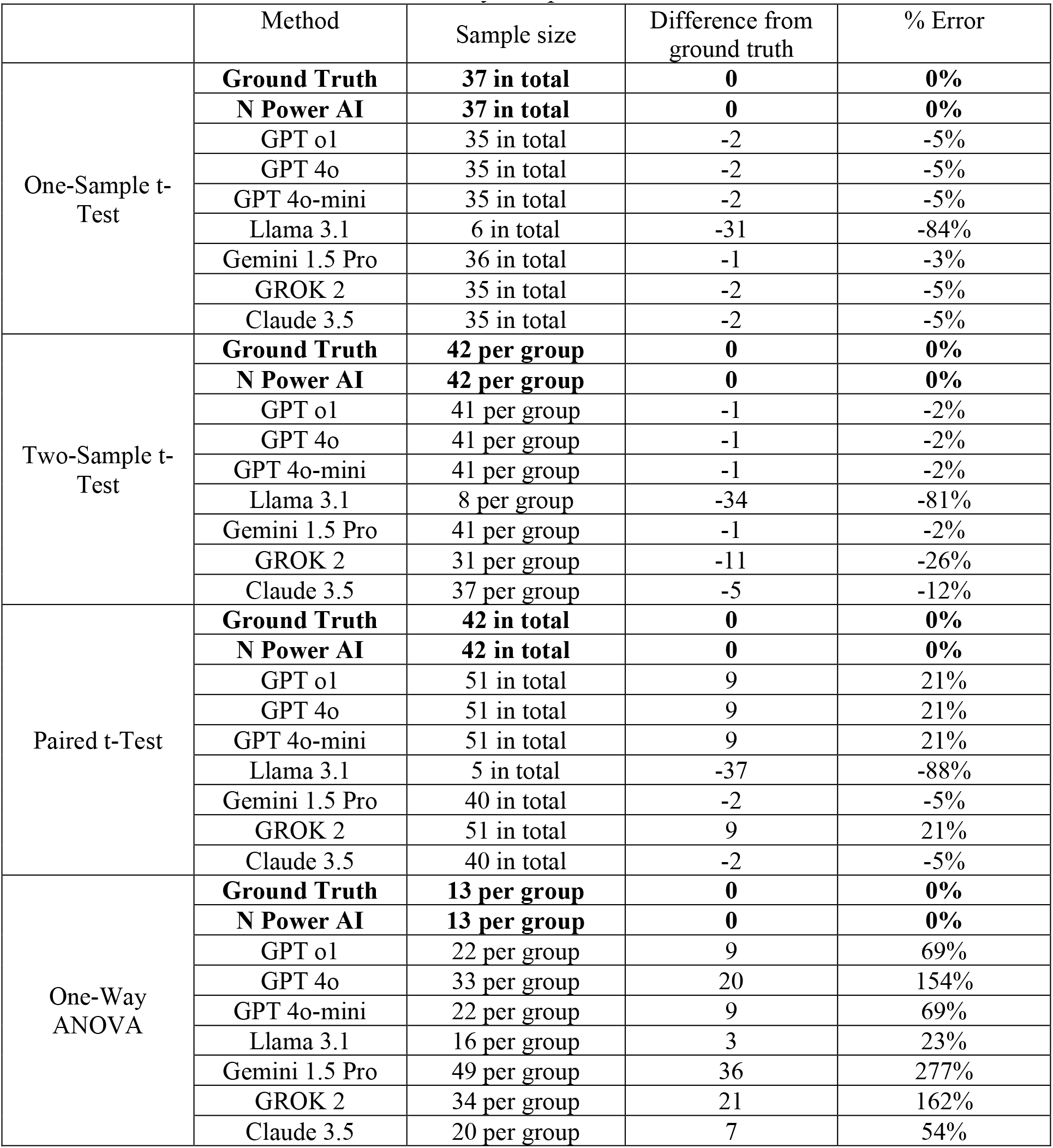
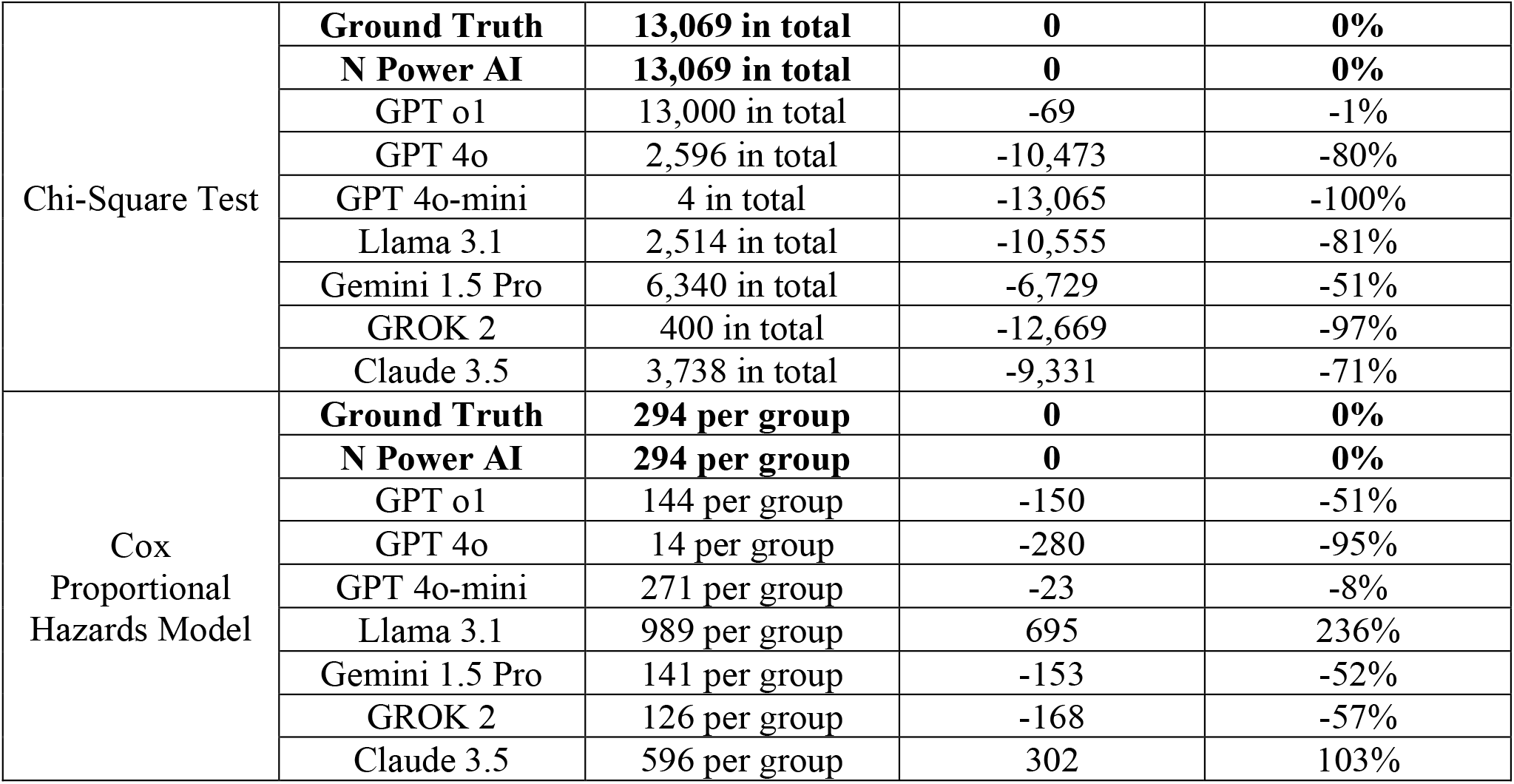
Sample size calculations across scenarios by the proposed N-Power AI and directly using various LLMs, including GPT-o1, GPT-4o, GPT-4o-mini, Llama 3.1, Claude 3.5, Gemini 1.5 Pro, and GROK 2, compared against the ground truth values derived from standard statistical software, R. Sample sizes are reported either as total values or per group where applicable. The proposed N-Power AI reported identical sample sizes in all ten replications, achieving 100% accuracy in sample size calculation across all six scenarios. In contrast, deviations from the ground truth were observed in sample sizes calculated by directly using various LLMs, particularly for more complex statistical models such as the Chi-Square Test and Cox Proportional Hazards Model, highlighting the limitations of direct LLM calls in handling intricate calculations and the necessity of structured frameworks of N-Power AI to ensure accuracy and precision in statistical tasks.

|  | Method | Sample size | Difference from ground truth | % Error |
| --- | --- | --- | --- | --- |
| One-Sample t-Test | **Ground Truth** | **37 in total** | **0** | **0%** |
|  | **N Power AI** | **37 in total** | **0** | **0%** |
|  | GPT o1 | 35 in total | -2 | -5% |
|  | GPT 4o | 35 in total | -2 | -5% |
|  | GPT 4o-mini | 35 in total | -2 | -5% |
|  | Llama 3.1 | 6 in total | -31 | -84% |
|  | Gemini 1.5 Pro | 36 in total | -1 | -3% |
|  | GROK 2 | 35 in total | -2 | -5% |
|  | Claude 3.5 | 35 in total | -2 | -5% |
| Two-Sample t-Test | **Ground Truth** | **42 per group** | **0** | **0%** |
|  | **N Power AI** | **42 per group** | **0** | **0%** |
|  | GPT o1 | 41 per group | -1 | -2% |
|  | GPT 4o | 41 per group | -1 | -2% |
|  | GPT 4o-mini | 41 per group | -1 | -2% |
|  | Llama 3.1 | 8 per group | -34 | -81% |
|  | Gemini 1.5 Pro | 41 per group | -1 | -2% |
|  | GROK 2 | 31 per group | -11 | -26% |
|  | Claude 3.5 | 37 per group | -5 | -12% |
| Paired t-Test | **Ground Truth** | **42 in total** | **0** | **0%** |
|  | **N Power AI** | **42 in total** | **0** | **0%** |
|  | GPT o1 | 51 in total | 9 | 21% |
|  | GPT 4o | 51 in total | 9 | 21% |
|  | GPT 4o-mini | 51 in total | 9 | 21% |
|  | Llama 3.1 | 5 in total | -37 | -88% |
|  | Gemini 1.5 Pro | 40 in total | -2 | -5% |
|  | GROK 2 | 51 in total | 9 | 21% |
|  | Claude 3.5 | 40 in total | -2 | -5% |
| One-Way ANOVA | **Ground Truth** | **13 per group** | **0** | **0%** |
|  | **N Power AI** | **13 per group** | **0** | **0%** |
|  | GPT o1 | 22 per group | 9 | 69% |
|  | GPT 4o | 33 per group | 20 | 154% |
|  | GPT 4o-mini | 22 per group | 9 | 69% |
|  | Llama 3.1 | 16 per group | 3 | 23% |
|  | Gemini 1.5 Pro | 49 per group | 36 | 277% |
|  | GROK 2 | 34 per group | 21 | 162% |
|  | Claude 3.5 | 20 per group | 7 | 54% |
| Chi-Square Test | **Ground Truth** | **13,069 in total** | **0** | **0%** |
|  | **N Power AI** | **13,069 in total** | **0** | **0%** |
|  | GPT o1 | 13,000 in total | -69 | -1% |
|  | GPT 4o | 2,596 in total | -10,473 | -80% |
|  | GPT 4o-mini | 4 in total | -13,065 | -100% |
|  | Llama 3.1 | 2,514 in total | -10,555 | -81% |
|  | Gemini 1.5 Pro | 6,340 in total | -6,729 | -51% |
|  | GROK 2 | 400 in total | -12,669 | -97% |
|  | Claude 3.5 | 3,738 in total | -9,331 | -71% |
| Cox<br>Proportional<br>Hazards Model | **Ground Truth** | **294 per group** | **0** | **0%** |
|  | **N Power AI** | **294 per group** | **0** | **0%** |
|  | GPT o1 | 144 per group | -150 | -51% |
|  | GPT 4o | 14 per group | -280 | -95% |
|  | GPT 4o-mini | 271 per group | -23 | -8% |
|  | Llama 3.1 | 989 per group | 695 | 236% |
|  | Gemini 1.5 Pro | 141 per group | -153 | -52% |
|  | GROK 2 | 126 per group | -168 | -57% |
|  | Claude 3.5 | 596 per group | 302 | 103% |

Table 2 shows similar results for power estimates as sample size calculation: none of the seven LLMs could return correct power estimates compared to the ground truth values derived from R crossing all six scenarios, particularly when more complex statistical models, like the Chi-Square Test and Cox Proportional Hazards Model, were needed. Furthermore, Gemini 1.5 Pro could not calculate precise power when the Chi-Square Test and Cox Proportional Hazards Model were needed, and it recommended users utilize software packages or online calculators to determine the specific power value for the study design. The above results suggest that directly asking LLMs to perform these calculations may lack the precision required for complex mathematical computations^30^. In addition, LLMs like GPT-4o tried to write and execute Python code to calculate the sample size for some scenarios. However, they still returned inaccurate sample sizes and power estimates, highlighting a critical limitation in their ability to perform statistical computations.

**Table 2.** Power estimates across scenarios by the proposed N-Power AI and directly using various LLMs, including GPT-o1, GPT-4o, GPT-4o-mini, Llama 3.1, Claude 3.5, Gemini 1.5 Pro, and GROK 2, compared against the ground truth values derived from standard statistical software, R. The proposed N-Power AI produced identical power estimations in all ten replications, achieving 100% accuracy in power estimation across all six scenarios. In contrast, deviations from the ground truth were observed in power estimates calculated by directly using various LLMs, particularly for more complex statistical models such as the Chi-Square Test and Cox Proportional Hazards Model. Gemini 1.5 Pro recommended that users utilize software packages or online calculators to determine the specific power value for the study design. These discrepancies highlight the limitations of direct LLM calls in handling intricate calculations and underscore the necessity of structured frameworks like N-Power AI to ensure accuracy and precision in statistical tasks.

|  | Method | Power estimates | Difference from ground truth | % Error |
| --- | --- | --- | --- | --- |
| One-Sample t-Test | **Ground Truth** | **0.9** | **0** | **0%** |
|  | **N Power AI** | **0.9** | **0** | **0%** |
|  | GPT o1 | 0.91 | 0.01 | 1% |
|  | GPT 4o | 0.085 | -0.815 | -91% |
|  | GPT 4o-mini | 0.92 | 0.02 | 2% |
|  | Llama 3.1 | 0.99 | 0.09 | 10% |
|  | Gemini 1.5 Pro | 0.91 | 0.01 | 1% |
|  | GROK 2 | 0.92 | 0.02 | 2% |
|  | Claude 3.5 | 0.93 | 0.03 | 3% |
| Two-Sample t-Test | **Ground Truth** | **0.81** | **0** | **0%** |
|  | **N Power AI** | **0.81** | **0** | **0%** |
|  | GPT o1 | 0.65 | -0.16 | -20% |
|  | GPT 4o | 0.88 | 0.07 | 9% |
|  | GPT 4o-mini | 0.65 | -0.16 | -20% |
|  | Llama 3.1 | 0.98 | 0.17 | 21% |
|  | Gemini 1.5 Pro | 0.99 | 0.18 | 22% |
|  | GROK 2 | 0.82 | 0.01 | 1% |
|  | Claude 3.5 | 0.85 | 0.04 | 5% |
| Paired t-Test | **Ground Truth** | **0.81** | **0** | **0%** |
|  | **N Power AI** | **0.81** | **0** | **0%** |
|  | GPT o1 | 0.71 | -0.1 | -12% |
|  | GPT 4o | 0.7 | -0.11 | -14% |
|  | GPT 4o-mini | 0.055 | -0.755 | -93% |
|  | Llama 3.1 | 0.99 | 0.18 | 22% |
|  | Gemini 1.5 Pro | 0.99 | 0.18 | 22% |
|  | GROK 2 | 0.92 | 0.11 | 14% |
|  | Claude 3.5 | 0.82 | 0.01 | 1% |
| One-Way ANOVA | **Ground Truth** | **0.81** | **0** | **0%** |
|  | **N Power AI** | **0.81** | **0** | **0%** |
|  | GPT o1 | 0.7 | -0.11 | -14% |
|  | GPT 4o | 0.69 | -0.12 | -15% |
|  | GPT 4o-mini | 0.92 | 0.11 | 14% |
|  | Llama 3.1 | 0.95 | 0.14 | 17% |
|  | Gemini 1.5 Pro | 0.63 | -0.18 | -22% |
|  | GROK 2 | 0.83 | 0.02 | 2% |
|  | Claude 3.5 | 0.83 | 0.02 | 2% |
| Chi-Square Test | **Ground Truth** | **0.8** | **0** | **0%** |
|  | **N Power AI** | **0.8** | **0** | **0%** |
|  | GPT o1 | 0.96 | 0.16 | 20% |
|  | GPT 4o | 1 | 0.2 | 25% |
|  | GPT 4o-mini | 1 | 0.2 | 25% |
|  | Llama 3.1 | 0.99 | 0.19 | 24% |
|  | Gemini 1.5 Pro | NA<sup>1</sup> | NA | NA |
|  | GROK 2 | 1 | 0.2 | 25% |
|  | Claude 3.5 | 0.95 | 0.15 | 19% |
| Cox Proportional Hazards Model | **Ground Truth** | **0.8** | **0** | **0%** |
|  | **N Power AI** | **0.8** | **0** | **0%** |
|  | GPT o1 | 0.86 | 0.06 | 7% |
|  | GPT 4o | 0.43 | -0.37 | -46% |
|  | GPT 4o-mini | 0.98 | 0.18 | 23% |
|  | Llama 3.1 | 1 | 0.2 | 25% |
|  | Gemini 1.5 Pro | NA | NA | NA |
|  | GROK 2 | 0.99 | 0.19 | 24% |
|  | Claude 3.5 | 0.89 | 0.09 | 11% |

<sup>1</sup>Not applicable. Gemini 1.5 Pro was unable to calculate the power estimates and recommended that users utilize software packages or online calculators to determine the specific power value for the study design.

In contrast to directly calling LLMs, the proposed N-Power AI framework, which integrates LLMs with specialized agents, achieved 100% accuracy in sample size calculation and power estimation across all six scenarios in all ten replications (Tables 1 and 2). These results highlight the necessity of an agentic framework to ensure accurate and reliable sample size and power calculations.

### PERFORMANCE OF THE THREE AGENTS IN THE N-POWER AI FRAMEWORK

We assess the performance of the three agents of N-Power AI. Table S5 presents the performance of the three agents of N-Power AI when various LLMs were chosen as the agents’ LLM settings. First of all, for the Function Agent, we examined whether it could identify suitable statistical models and select the correct R functions from the predefined R function repository for downstream agents. Table S5 shows that the Function Agent achieved 100% accuracy in both identifying appropriate statistical models and selecting the corresponding R functions across different LLM settings. This finding suggests the robustness of the Function Agent in interpreting study descriptions and aligning them with the correct statistical methodologies, which is a critical step in ensuring the reliability of the N-Power AI framework. Next, we assessed the Calculation Agent’s performance, concentrating on two main tasks: (1) accurately extracting the necessary parameters for the selected R function from the input scenarios and (2) correctly calculating sample sizes and power estimates. Again, the Calculation Agent achieved 100% accuracy in extracting the required parameters, such as effect size, significance level, and power, ensuring that the inputs fed into the R functions are both complete and valid across different LLM settings (Table S5). The Calculation Agent also achieved 100% accuracy in calculating sample sizes and power estimates across different LLM settings (Table S5), highlighting the agent’s ability to execute complex statistical calculations reliably, consistently aligning with the ground-truth results generated by R.

Finally, we evaluated the performance of the Reporting Agent by manually examining whether the generated reports accurately reflected the study objectives, hypotheses, selected statistical models, underlying assumptions, and detailed analysis results. The Reporting Agent achieved 100% accuracy in compiling these elements into a comprehensive and user-friendly report across different LLM settings (Table S5), which ensures that the final output is not only statistically rigorous but also accessible to researchers with varying levels of statistical expertise. We chose Llama 3.1 70b-instruct-8k as the LLM model for N-Power AI due to its open-access availability, performance, and cost-free usage, making it an ideal choice for researchers seeking an accessible and reliable solution.

### N-POWER AI GENERATES COMPREHENSIVE REPORTS

N-Power AI generates comprehensive, user-friendly reports that summarize the entire analysis process, making it an invaluable tool for researchers. The Reporting Agent compiles detailed summaries that include study objectives, hypotheses, selected statistical models, underlying assumptions, and precise analysis results (Figure 2, Figure S1-S5). These reports are intended to be informative and accessible, serving researchers with different levels of statistical expertise.

## Discussion

The development and evaluation of N-Power AI demonstrate the potential of leveraging LLMs within a specialized agent framework to automate statistical planning for clinical trial design. Our results show that N-Power AI achieves 100% accuracy in identifying appropriate statistical models, selecting correct R functions, extracting parameters, calculating sample sizes and power estimates, and generating comprehensive reports across six common clinical trial scenarios, highlighting the accuracy and robustness of the framework. Importantly, the three-LLM-based agent architecture— comprising the Function Agent, Calculation Agent, and Reporting Agent—ensures that each step of the process is handled with domain-specific expertise and computational rigor, addressing the limitations of directly using LLM.

The success of N-Power AI lies in its ability to combine the natural language processing strengths of LLMs with the precision of statistical software through the function calling approach. Directly using LLMs alone lacks the ability to perform precise mathematical computations. While ChatGPT can generate and execute Python code, they still often produce inaccurate results. This limitation underscores the necessity of a structured, agent-based approach. By integrating LLMs with specialized agents, N-Power AI not only automates complex statistical tasks but also ensures that the outputs are reliable, reproducible, and aligned with best practices in clinical trial design. The framework’s ability to generate comprehensive, user-friendly reports further enhances its utility. Researchers can easily interpret and share the results, making N-Power AI a valuable tool for clinical trial design, grant applications, study protocols, and academic publications. Moreover, the open-access nature of the Llama 3.1 70b model used in the framework ensures that the tool is accessible to a wide range of researchers.

While N-Power AI demonstrates significant promise, several limitations should be acknowledged. First, the scenarios tested in this study are limited to six common clinical trial designs. While these scenarios cover a broad range of applications, they do not include all possible study designs or statistical methods. Future work should expand the evaluation to include more complex scenarios to further validate the framework’s generalizability. Second, the framework relies on predefined R functions from packages like pwrss, which may not cover all statistical methods or study designs. Incorporating additional statistical packages, custom R scripts or even functions generated by specialized coding LLMs (i.e., Code Llama) could enhance the framework’s versatility. Finally, the framework’s reliance on LLMs introduces potential biases or limitations inherent to these LLMs, such as sensitivity to prompt phrasing. While the specialized agent framework mitigates many of these issues, ongoing monitoring and updates will be necessary to ensure that N-Power AI remains aligned with evolving best practices in both AI and clinical trial design.

Despite these limitations, N-Power AI represents a significant step forward in statistical planning for clinical trials. By combining the strengths of LLMs with an agent-based framework, the tool makes statistical planning more accessible, efficient, and reliable. While human expertise remains essential for ensuring the scientific rigor of clinical research, N-Power AI has the potential to provide access to advanced statistical tools, particularly for researchers with limited resources or expertise. N-Power AI is freely available at https://ai.swmed.edu/N-PowerAI/.

## Notes

### Competing Interest Statement

The authors have declared no competing interest.

## Reference

1. Piantadosi S. Clinical trials: a methodologic perspective. John Wiley & Sons, 2024.

2. Chow SC, Liu JP. Design and analysis of clinical trials: concepts and methodologies. John Wiley & Sons, 2008.

3. R Core Team. R: A language and environment for statistical computing. R Foundation for Statistical Computing. 2021 (http://www.R-project.org/).

4. SAS Institute Inc. Base SAS 9.4 Procedures Guide. 2015 (http://support.sas.com).

5. Faul F, Erdfelder E, Lang AG, Buchner A. G* Power 3: A flexible statistical power analysis program for the social, behavioral, and biomedical sciences. Behavior Research Methods. 2007;39:175–91.

6. Thirunavukarasu AJ, Ting DS, Elangovan K, Gutierrez L, Tan TF, Ting DS. Large language models in medicine. Nature Medicine. 2023;29(8):1930–40.

7. Jiang J, Wang F, Shen J, Kim S, Kim S. A Survey on Large Language Models for Code Generation. 2024. (https://arxiv.org/abs/2406.00515). Preprint.

8. Jin Q, Wang Z, Floudas CS, Chen F, Gong C, Bracken-Clarke D, Xue E, Yang Y, Sun J, Lu Z. Matching patients to clinical trials with large language models. Nature Communications. 2024;15:9074.

9. Fast D, Adams LC, Busch F, Fallon C, Huppertz M, Siepmann R, Prucker P, Bayerl N, Truhn D, Makowski M, Löser A. Autonomous medical evaluation for guideline adherence of large language models. NPJ Digital Medicine. 2024;7:1–4.

10. Ayers JW, Poliak A, Dredze M, Leas EC, Zhu Z, Kelley JB, Faix DJ, Goodman AM, Longhurst CA, Hogarth M, Smith DM. Comparing physician and artificial intelligence chatbot responses to patient questions posted to a public social media forum. JAMA Internal Medicine. 2023;183:589–96.

11. Durante Z, Huang Q, Wake N, Gong R, Park JS, Sarkar B, Taori R, Noda Y, Terzopoulos D, Choi Y, Ikeuchi K. Agent AI: Surveying the horizons of multimodal interaction. 2024. (https://arxiv.org/abs/2401.03568). Preprint.

12. Qiu, J., Lam, K., Li, G., Acharya, A., Wong, T. Y., Darzi, A., … & Topol, E. J. (2024). LLM-based agentic systems in medicine and healthcare. Nature Machine Intelligence, 6(12), 1418–1420.

13. Anthropic AI. Building effective agents. 2024. (https://www.anthropic.com/research/building-effective-agents).

14. Laurent G, Michał K. rpy2 - R in Python. Oct. 03, 2024. (https://rpy2.github.io/).

15. Ollama. accessed Oct. 03, 2024. (https://ollama.com/)

16. META. Introducing Llama 3.1: Our most capable models to date. July 23, 2024. (https://ai.meta.com/blog/meta-llama-3-1/).

17. OpenAI. API platform. accessed Oct. 03, 2024. (https://openai.com/api/).

18. Bulus, M. pwrss: Statistical Power and Sample Size Calculation Tools. R package version 0.3.1. 2023. (https://CRAN.R-project.org/package=pwrss).

19. Basu, K. (2024, October). Bridging Knowledge Gaps in LLMs via Function Calls. In Proceedings of the 33rd ACM International Conference on Information and Knowledge Management (pp. 5556–5557).

20. Daniel WW, Cross CL. Biostatistics: a foundation for analysis in the health sciences. Wiley; Nov 13, 2018.

21. Rosner BA. Fundamentals of biostatistics. Boston, MA: Brooks/Cole, Cengage Learning; 2015.

22. Gao W, Ping S, Liu X. Gender differences in depression, anxiety, and stress among college students: a longitudinal study from China. Journal of Affective Disorders. Feb 15, 2020;263:292–300.

23. UCLA: Statistical Consulting Group. Data Analysis Examples. Accessed Oct. 03, 2024. (https://stats.oarc.ucla.edu/other/dae/)

24. OpenAI. Introducing OpenAI o1-preview. September 12, 2024. (https://openai.com/index/introducing-openai-o1-preview/).

25. OpenAI. Hello GPT-4o. May 13, 2024. (https://openai.com/index/hello-gpt-4o/).

26. OpenAI. GPT-4o mini: advancing cost-efficient intelligence. July 18, 2024. (https://openai.com/index/gpt-4o-mini-advancing-cost-efficient-intelligence/).

27. Anthropic. Claude 3.5 Sonnet. June 20, 2024. (https://www.anthropic.com/news/claude-3-5-sonnet).

28. Team G, Georgiev P, Lei VI, Burnell R, Bai L, Gulati A, Tanzer G, Vincent D, Pan Z, Wang S, Mariooryad S. Gemini 1.5: Unlocking multimodal understanding across millions of tokens of context. Mar 8, 2024. (https://arxiv.org/abs/2403.05530). Preprint.

29. X. Bringing Grok to Everyone. December 12, 2024. (https://x.ai/blog/grok-1212).

30. Collins KM, Jiang AQ, Frieder S, Wong L, Zilka M, Bhatt U, Lukasiewicz T, Wu Y, Tenenbaum JB, Hart W, Gowers T. Evaluating language models for mathematics through interactions. Proceedings of the National Academy of Sciences. Jun 11, 2024;121(24):e2318124121.

