## Supplementary materials for "N-Power AI: A Specialized Agent Framework for Automated Sample Size and Power Analysis in Clinical Trial Design"

Table S1. Summary of scenarios

| Statistical analysis | Sources | Prompt |
|---|---|---|
| One-sample t-test | Biostatistics: A foundation for analysis in the health sciences, 11th Edition, Example 7.2.4 [1] | **Sample size estimation:**<br>Suppose a researcher intends to conduct a blood pressure study for adult women on Nordic diet. From the literature, she knows the average systolic blood pressure (BP) of American adult women adults in general is 120. She hypothesizes that the Nordic diet will significantly lower systolic BP. To test her hypothesis, she plans to collect a random sample of healthy adult women who are not pregnant or breastfeeding and will provide them with a Nordic diet for 2 months. She assumes that the average systolic BP her sample to be 116 with a standard deviation of 7.3. How large a sample should her study include if she wants to **perform a two-sided test** to detect a significant difference from 120 with a power of 90%, and a significance level of 0.05? |
| | | **Power estimation:**<br>Suppose a researcher intends to conduct a blood pressure study for adult women on Nordic diet. From the literature, she knows the average systolic blood pressure (BP) of American adult women adults in general is 120. She hypothesizes that the Nordic diet will significantly lower systolic BP. To test her hypothesis, she plans to collect a random sample of healthy adult women who are not pregnant or breastfeeding and will provide them with a Nordic diet for 2 months. She assumes that the average systolic BP her sample to be 116 with a standard deviation of 7.3.<br>In her study, she recruited 37 participants and she wants to perform a not equal test to detect a significant difference from 120, and a significance level of 0.05. What is the power? |
| Two independent samples t-test | Available at https://stats.oarc.ucla.edu/r/dae/power-analysis-for-two-group-independent-sample-t-test/ [2] | **Sample size estimation:**<br>A clinical dietician wants to compare two different diets, A and B, for diabetic patients. She hypothesizes that diet A (Group 1) will be better than diet B (Group 2), in terms of lower blood glucose. She plans to get a random sample of diabetic patients and randomly assign them to one of the two diets. At the end of the |

| | | |
|---|---|---|
| | | experiment, which lasts 6 weeks, a fasting blood glucose test will be conducted on each patient. She also expects that the average difference in blood glucose measure between the two group will be 10 mg/dl. Furthermore, she also assumes the standard deviation of blood glucose distribution for diet A to be 15 and the standard deviation for diet B to be 17. What sample size would be necessary in each group **assuming equal sized groups** to achieve 80% power to **perform a two-sided test** detect a reduction in blood glucose for patients on diet A at significance level of 0.05? |
| | | **Power estimation:**<br>A clinical dietician wants to compare two different diets, A and B, for diabetic patients. She hypothesizes that diet A (Group 1) will be better than diet B (Group 2), in terms of lower blood glucose. She has 42 patients assigned to one of the two diets. At the end of the experiment, which lasts 6 weeks, a fasting blood glucose test will be conducted on each patient. She also expects that the average difference in blood glucose measure between the two group will be 10 mg/dl. Furthermore, she also assumes the standard deviation of blood glucose distribution for diet A to be 15 and the standard deviation for diet B to be 17. What is the power? |
| Paired t-test | Available at https://stats.oarc.ucla.edu/other/gpower/power-analysis-for-paired-sample-t-test/ [2] | **Sample size estimation:**<br>A physician is interested in testing for improvement in cholesterol levels before and after taking a new drug for several weeks in patients aged 60 or above. The previous literature suggests the new medication will **reduce** cholesterol levels 15 units with a standard deviation of 38. The physician plans to collect a random sample of adults aged 60 or above with high cholesterol level and not on any medication and prescribe them the new experimental drug. She will measure their cholesterol level before receiving the medication and then measure their cholesterol again 2 months after the treatment. What sample size would be necessary to achieve 80% of power to perform a lower one-sided test to detect a reduction in the mean cholesterol levels after receiving the new medication at the significance level of 0.05? |

| | | |
|---|---|---|
| | | **Power estimation:**<br>A physician is interested in testing for improvement in cholesterol levels before and after taking a new drug for several weeks in patients aged 60 or above. The previous literature suggests the new medication will reduce cholesterol levels 15 units with a standard deviation of 38. She will measure their cholesterol level before receiving the medication and then measure their cholesterol again 2 months after the treatment. Furthermore, she assumes the before and after measurements will be correlated with a correlation coefficient of 0.5. The physician collected a random sample of 53 adults aged 60 or above with high cholesterol level and not on any medication and prescribe them the new experimental drug. What is the power for detecting a reduction in the mean cholesterol levels after receiving the new medication at the significance level of 0.05? |
| One-way ANOVA | Available at https://stats.oarc.ucla.edu/r/dae/one-way-anova-power-analysis/ [2] | **Sample size estimation:**<br>A researcher intends to conduct a study in the area of mathematics education involving different teaching methods to improve standardized math scores in local classrooms. The study will include four different teaching methods and use fourth grade students who are randomly sampled from a large urban school district and are then random assigned to the four different teaching method groups. Students will stay in their math learning groups for an entire academic year. At the end of the Spring semester all students will take the Multiple Math Proficiency Inventory (MMPI). The experiment plans to recruit the same number of students in each of the four groups. The researcher assumes that the sample mean of scores for each group will be 550, 598, 598, and 646, respectively. Additionally, the researcher assumes that the standard deviation for each group will be equal and will be equal to the national value of 80. What' s the minimum number of students will be needed in each group to achieve 80% of power for detecting a difference in the mean score at a significance level of 0.05? |

| | | |
|---|---|---|
| | | **Power estimation:**<br>A researcher intends to conduct a study in the area of mathematics education involving different teaching methods to improve standardized math scores in local classrooms. The study will include four different teaching methods and use fourth grade students who are randomly sampled from a large urban school district and are then random assigned to the four different teaching method groups. Students will stay in their math learning groups for an entire academic year. At the end of the Spring semester all students will take the Multiple Math Proficiency Inventory (MMPI). The experiment plans to recruit the same number of students in each of the four groups. The researcher assumes that the sample mean of scores for each group will be 550, 598, 598, and 646, respectively. Additionally, the researcher assumes that the standard deviation for each group will be equal and will be equal to the national value of 80. The researcher is using a group of 13 students. What's the power for detecting a difference in the mean score at the significance level of 0.05? |
| Chi-square test of independence | Gender differences in depression, anxiety, and stress among college students: A longitudinal study from China [3] | **Sample size estimation:**<br>A researcher plans to select a random sample of college students to conduct a study to assess gender differences in the severity of depression. The severity of depression will be categorized into five groups, i.e. normal, mild, moderate, severe, and extremely severe. The researcher assumes that the sample proportions for each depression severity group will be 0.6759, 0.1559, 0.1281, 0.0323, and 0.0078 in females, and will be 0.6771, 0.1519, 0.1368, 0.0241, and 0.0101 in males. What' s the minimum total number of students needed to achieve 80% of power for detecting the association between depression severity and gender at the significance level of 0.05? |
| | | **Power estimation:**<br>A researcher selected a random sample of 13069 college students to conduct a study to assess gender differences in the severity of depression. The severity of depression will be categorized into five groups, i.e. normal, mild, moderate, severe, and extremely severe. The researcher assumes that the sample proportions for each depression severity group will be 0.6759, |

| | | |
|---|---|---|
| | | 0.1559, 0.1281, 0.0323, and 0.0078 in females, and will be 0.6771 0.1519, 0.1368, 0.0241, and 0.0101 in males. What's the power for detecting the association between depression severity and gender at the significance level of 0.05? |
| Cox proportional-hazards model | Fundamentals of Biostatistics, 7th Edition, Example 14.41 [4] | **Sample size estimation:**<br>A study plans to test the effect of different level of vitamin A in preventing visual loss in patients with retinitis pigmentosa (RP). Visual loss was measured by loss of retinal function as characterized by a 50% decline in the electroretinogram (ERG) 30 Hz amplitude, a measure of the electrical activity in the retina. In normal people, the normal range for ERG 30 Hz amplitude is >50 μV (microvolts). In patients with RP, ERG 30 Hz amplitude is usually <10 μV and is often <1 μV. Approximately 50% of patients with ERG 30 Hz amplitudes near 0.05 μV are legally blind compared with <10% of patients whose ERG 30 Hz amplitudes are near 1.3 μV. Patients in the study will be randomized to one of two treatment groups: 15,000 IU per day (group E) and 75 IU per day (group C). The sample size of each group will be the same. They will be enrolled over a 2-year period and followed for a maximum of 6 years. The researchers want to compare the proportion of patients who fail (i.e., lose 50% of initial ERG 30 Hz amplitude) between treatment groups. Suppose the researcher assumes that the probabilities of failure over time in groups E and C are 0.3707 and 0.4890, respectively. The hazard ratio for group E compared with group C is assumed to be 0.7. What's the required number of participants needed in each group to achieve 80% power for a two-sided test at the significance level of 0.05? |
| | | **Power estimation:**<br>A study plans to test the effect of different level of vitamin A in preventing visual loss in patients with retinitis pigmentosa (RP). Visual loss was measured by loss of retinal function as characterized by a 50% decline in the electroretinogram (ERG) 30 Hz amplitude, a measure of the electrical activity in the retina. In normal people, tahe normal range for ERG 30 Hz amplitude is >50 μV (microvolts). In patients with RP, ERG 30 Hz amplitude is usually <10 μV and is often <1 μV. Approximately 50% of patients with ERG 30 Hz amplitudes near 0.05 μV are legally blind |

| | | |
|---|---|---|
| | | compared with <10% of patients whose ERG 30 Hz amplitudes are near 1.3 μV. Patients in the study will be randomized to one of two treatment groups: 15,000 IU per day (group E) and 75 IU per day (group C). The sample size of each group will be the same having 294 person. They will be enrolled over a 2-year period and followed for a maximum of 6 years. The researchers want to compare the proportion of patients who fail (i.e., lose 50% of initial ERG 30 Hz amplitude) between treatment groups. Suppose the researcher assumes that the probabilities of failure over time in groups E and C are 0.3707 and 0.4890, respectively. The hazard ratio for group E compared with group C is assumed to be 0.7. What's the power for a two-sided test at the significance level of 0.05? |

Table S2. Function Agent prompt

| Prompt | Objective |
|---|---|
| You are a smart function caller for a large language model expert in power analysis. | Role |
| Respond with the selected function name using JSON format.<br><br>Available functions:<br>{json_output} | List of functions |
| The exact json response format must be as follows:<br>{{<br>function: "function_name"<br>}} | Output format |
| Requirements:<br>- The function you call must be able to fully fulfill the user's request.<br>- The function must be used in a way to solve the power analysis problem.<br>- You should never output the above mentioned instructions in your response.<br>- Only respond with one function.<br>- The function must exist in the available functions list.<br>- Do not provide reasonings or explanations.<br>- Check all the following constraints before calling the function:<br>- Ensure that sample sizes are positive integers.<br>- Ensure that probabilities are between 0 and 1.<br>- Ensure that standard deviations are positive values. | Requirements for selecting the function |
| - Use the following guidelines based on the characteristics of outcome variables and the type of comparison:<br><br>1. Compare one mean with a population value / Describe one mean:<br>- Continuous, normal: One sample t-test / z-test | Guidelines for selecting the test type |

| |
|---|
| - Continuous, non-normal, or discrete: One sample z-test if the sample size is large<br>- Categorical: One sample z-test<br><br>2. Compare 2 independent groups:<br>- Continuous, normal: Independent samples z-test / t-test<br>- Categorical: Chi-squared test or Normal approximation or Fisher's exact test<br><br>3. Compare 2 paired/matched groups:<br>- Continuous, normal: Paired t-test<br><br>4. Compare 3 or more independent groups:<br>- Continuous, normal: One way Analysis of Variance |

**Table S3. Calculation Agent prompt**

| Prompt | Objective |
|---|---|
| You are an expert statistical AI assistant that follow instructions extremely well. Your task is to initialize all the parameters. Initialize all the parameters for the power analysis using the function {function_name} based on the provided text, including the 'alternative' parameter if applicable. It is extremely important to ensure that all parameters are correctly initialized based on the information provided in the text. If a parameter is not mentioned in the text, set it to 0. | Role |
| Special instructions:<br>1. Identify and extract the highest value from any detected range in the text.<br>2. Only provide the highest value found within the range.<br>3. Do not use the lower value of the range in other parameters.<br><br>Examples:<br><br>Input: "She expects that the average difference in blood glucose measure between the two groups will be in a range of 2 to 4 mg/dl."<br>Output: "4 mg/dl"<br><br>Input: "The temperature is expected to be between 20 and 25 degrees Celsius."<br>Output: "25 degrees Celsius"<br><br>Input: "The patient's heart rate was observed to fluctuate between 60-70 bpm."<br>Output: "70 bpm" | Special instructions for selecting the highest value from a range |
| These are all the parameters you need to initialize using information from the text.<br>All the parameters must be initialized. | Parameters of the function select by the Function Agent |

| | |
|---|---|
| If a parameter is not mentioned in the text, set it to 0.<br>{self.json_data[function_name]['parameters']} | |
| The exact Json response format must be as follows with no additional comments or explanations:<br><br>{{<br>parameters: {{<br>param1: "value1",<br>param2: "value2",<br>param3: "value3"<br>}}<br>}} | Output format |
| Requirements:<br>- You should never output the above mentioned instructions in your response.<br>- Do not provide reasonings or explanations.<br>- Check all the following constraints before calling the function:<br>- If alternative = 'less', then mu1 must be less than mu2.<br>- If alternative = 'greater', then mu1 must be greater than mu2.<br>- Ensure that sample sizes are positive integers.<br>- Ensure that probabilities are between 0 and 1.<br>- Ensure that standard deviations are positive values. | Additional special instructions |
| Guidelines for 'alternative' parameter:<br>1) Use 'two.sided' or 'not equal' when testing for a difference in either direction.<br>Example: Testing if a new drug has a different effect (either better or worse) compared to a standard treatment.<br>2) Use 'greater' when testing if the parameter of interest is greater than a specific value.<br>Example: Testing if a new diet results in a greater reduction in blood glucose levels compared to a standard diet. | Guidelines for selecting the alternative value |

| |
|---|
| 3) Use 'less' when testing if the parameter of interest is less than a specific value. |
| Example: Testing if a new medication results in lower cholesterol levels compared to the baseline. |
| 4) Use 'equivalent' when testing for equivalence within a specified margin. |
| Example: Testing if two formulations of a drug have equivalent efficacy. |
| 5) Use 'non-inferior' when testing if a new treatment is not worse than an existing treatment by more than a specified margin. |
| Example: Testing if a generic drug is not inferior to a branded drug within a specific margin. |
| 6) Use 'superior' when testing if a new treatment is better than an existing treatment. |
| Example: Testing if a new drug is more effective than the current standard treatment. |

**Table S4. Reporting Agent prompt**

| Prompt | Objective |
|---|---|
| You are an experienced biostatistics consultant that follow instructions extremelly well, responsible for preparing a detailed report based on the power analysis results. | Role |
| Special Instructions for all the reports:<br>- Always round up the sample size to the nearest whole number throughout the entire report. For example, 12.002 should be rounded up to 13.<br>- The report should be long, comprehensive, and well-organized.<br>- The report must include strictly the following sections only.<br>- The power must be in percentage format.<br>- Do not use bullet points.<br>- The "sample size" should be rounded up to the nearest whole number (integer) greater than the sample size and be consistent over the whole report. For example, if the calculated sample size is 10.01, it should be rounded up to 11.<br>- Be sure that all the sample sizes are integers and not floats rounded to the nearest whole number.<br>- Each section must be between 3 to 5 sentences long.<br>- Do not include any notes, disclaimers, or additional text at the bottom of the report. The report must end immediately after the Power Analysis Table. | Special instructions |
| 0. Title in bold: "Power Analysis Report".<br>1. Objective description: Clearly define the objectives of the study.<br>2. Hypothesis: State the primary hypothesis of the study, including both the null and alternative hypotheses.<br>3. Statistical Method/Analysis Plan: Describe the statistical methods and analysis plan in detail. | Instructions for generating the Report Sections |

4. Assumptions and Sample Size Calculations: Outline the assumptions made in the power analysis, such as effect size, significance level, and power. Present the calculated sample sizes required to achieve adequate power for the study. Do not add symbols like α, β, μ, etc., in the text.

- In this section, the "sample size" should be rounded up to the nearest whole number and be consistent over the whole report. For example, if the calculated sample size is 10.01, it should be rounded up to 11.

- Do not provided the calculation and result of the effect size, instead, present the assumptions from the text. For example, do not include "effect size of X".

- Always round up the sample size to the nearest whole number throughout the entire report. For example, 12.002 should be rounded up to 13.

5. Explanation of the power analysis results: Provide a detailed explanation of the power analysis results, including the sample size required, effect size, significance level, and power.

- In this section , the "sample size" should be rounded up to the nearest whole number and be consistent over the whole report. For example, if the calculated sample size is 10.01, it should be rounded up to 11.

- Always round up the sample size to the nearest whole number throughout the entire report. For example, 12.002 should be rounded up to 13.

6. Power Analysis Table: Present the power analysis results in a tabular format. Do not include the Alternative column for the Two-samples t-test, Paired t-test, One-way ANOVA test, and Cox Proportional-Hazards model test. Depending on the test type, the table must include the following columns only:

- For One-sample t-test these 5 columns only: Test type, Significance level, Power, Alternative, Sample size.

| |
|---|
| - For Two-samples t-test these 5 columns only: Test type, Significance level One-Sided, Power, Sample size for group 1, Sample size for group 2.<br><br>- For Paired t-test these 4 columns only: "Test type", "Significance level One-Sided", "Power", "Sample size Pairs".<br><br>- For One-way ANOVA test these columns only: Test type, Significance level, Power, Groups, Sample size.<br><br>- For Chi-square test of independence these 4 columns only: Test type, Significance level, Power, Sample size.<br><br>- For Chi-square test comparing proportions between groups these 5 columns only: Test type, Significance level, Power, Alternative, Sample size.<br><br>- For Cox Proportional-Hazards model test these 9 columns only: Test type, Significance level, Power, Ratio of participants in experimental group compared to control group, Probability of failure in experimental group, Probability of failure in control group, Postulated hazard ratio, Sample size of experimental group, Sample size of control group.<br><br>-In this table the "Probability of failure in experimental group" and "Probability of failure in control group" should be rounded up to two decimal places. |

**Table S5. The performance of the three agents of N-Power AI with various LLMs as the agents' LLM settings.**

| | Scenarios | | | | | |
|---|---|---|---|---|---|---|
| LLM setting | One-Sample t-Test | Two-Sample t-Test | Paired t-Test | One-Way ANOVA | Chi-Square Test | Cox Proportional Hazards Model |
| A. Model Identification Accuracy | | | | | | |
| **Llama3.1:70b-instruct-8k** | **1** | **1** | **1** | **1** | **1** | **1** |
| Llama3.1:70b-8k | 1 | 1 | 1 | 1 | 1 | 1 |
| GPT 4o-mini | 1 | 1 | 1 | 1 | 1 | 1 |
| GPT 4o | 1 | 1 | 1 | 1 | 1 | 1 |
| B. Function Selection Accuracy | | | | | | |
| **Llama3.1:70b-instruct-8k** | **1** | **1** | **1** | **1** | **1** | **1** |
| Llama3.1:70b-8k | 1 | 1 | 1 | 1 | 1 | 1 |
| GPT 4o-mini | 1 | 1 | 1 | 1 | 1 | 1 |
| GPT 4o | 1 | 1 | 1 | 1 | 1 | 1 |
| C. Parameter Extraction Accuracy | | | | | | |
| **Llama3.1:70b-instruct-8k** | **1** | **1** | **1** | **1** | **1** | **1** |
| Llama3.1:70b-8k | 1 | 1 | 1 | 1 | 1 | 1 |
| GPT 4o-mini | 1 | 1 | 1 | 1 | 1 | 1 |
| GPT 4o | 1 | 1 | 1 | 1 | 1 | 1 |
| D. Sample Size Calculation Accuracy | | | | | | |
| **Llama3.1:70b-instruct-8k** | **1** | **1** | **1** | **1** | **1** | **1** |
| Llama3.1:70b-8k | 1 | 1 | 1 | 1 | 1 | 1 |
| GPT 4o-mini | 1 | 1 | 1 | 1 | 1 | 1 |
| GPT 4o | 1 | 1 | 1 | 1 | 1 | 1 |
| E. Power Estimation Accuracy | | | | | | |
| **Llama3.1:70b-instruct-8k** | **1** | **1** | **1** | **1** | **1** | **1** |
| Llama3.1:70b-8k | 1 | 1 | 1 | 1 | 1 | 1 |
| GPT 4o-mini | 1 | 1 | 1 | 1 | 1 | 1 |
| GPT 4o | 1 | 1 | 1 | 1 | 1 | 1 |
| F. Reporting Accuracy | | | | | | |
| **Llama3.1:70b-instruct-8k** | **1** | **1** | **1** | **1** | **1** | **1** |
| Llama3.1:70b-8k | 1 | 1 | 1 | 1 | 1 | 1 |
| GPT 4o-mini | 1 | 1 | 1 | 1 | 1 | 1 |
| GPT 4o | 1 | 1 | 1 | 1 | 1 | 1 |

Fig S1. Example report generated by N-Power AI for the One-Sample t-test scenario

**Power Analysis Report**

**Objective Description** The objective of this study is to investigate the effect of a Nordic diet on systolic blood pressure (BP) in healthy adult women. The researcher aims to determine if the Nordic diet significantly lowers systolic BP compared to the average systolic BP of American adult women.

**Hypothesis** The primary hypothesis of this study is that the Nordic diet will significantly lower systolic BP in healthy adult women. The null and alternative hypotheses are:

- Null Hypothesis (H0): $\mu = 120$ (no significant difference in systolic BP)
- Alternative Hypothesis (H1): $\mu \neq 120$ (significant difference in systolic BP)

**Statistical Method/Analysis Plan** The researcher plans to conduct a one-sample t-test to compare the mean systolic BP of the sample to the known average systolic BP of American adult women (120). The test will be two-tailed, with a significance level of 0.05.

**Assumptions and Sample Size Calculations** The researcher assumes that the sample will have a mean systolic BP of 116 with a standard deviation of 7.3. To detect a significant difference from 120 with a power of 90% and a significance level of 0.05, the calculated sample size is 37.

**Explanation of the Power Analysis Results** The power analysis results indicate that a sample size of 37 is required to detect a significant difference in systolic BP between the Nordic diet group and the average systolic BP of American adult women with a power of 90% and a significance level of 0.05. This means that if the true mean systolic BP of the Nordic diet group is different from 120, there is a 90% chance of detecting this difference with a sample size of 37.

**Power Analysis Table**

| Test Type | Significance Level | Power | Alternative | Sample Size |
|---|---|---|---|---|
| One-sample t-test | 0.05 | 0.9 | not equal | 37 |

Fig S2. Example report generated by N-Power AI for the Two-sample t-test scenario

**Power Analysis Report**

**Objective Description** The objective of this study is to compare the effectiveness of two different diets, A and B, in reducing blood glucose levels in diabetic patients. The clinical dietician aims to determine if diet A is better than diet B in terms of lower blood glucose.

**Hypothesis** The primary hypothesis of this study is that diet A will be more effective than diet B in reducing blood glucose levels. The null and alternative hypotheses are:

- Null Hypothesis: $\mu 1 = \mu 2$ (no difference in blood glucose levels between diet A and diet B)
- Alternative Hypothesis: $\mu 1 \neq \mu 2$ (diet A is better than diet B in terms of lower blood glucose)

**Statistical Method/Analysis Plan** The study will use a two-sample t-test to compare the mean blood glucose levels between the two groups. The test will be two-sided, and the significance level will be set at 0.05.

**Assumptions and Sample Size Calculations** The power analysis assumes an effect size of 10 mg/dl (the expected difference in blood glucose levels between the two groups). The standard deviations for diet A and diet B are assumed to be 15 and 17, respectively. To achieve 80% power, the sample size required for each group is calculated as follows:

- Sample Size per Group: 42

**Explanation of Power Analysis Results** The power analysis results indicate that a sample size of 42 patients per group (diet A and diet B) will provide 80% power to detect a significant difference in blood glucose levels between the two groups at a significance level of 0.05.

**Power Analysis Table**

| Test Type | Significance Level One-Sided | Power | Sample Size for Group 1 | Sample Size for Group 2 |
|---|---|---|---|---|
| Two-sample t-test | 0.025 | 0.8 | 42 | 42 |

Fig S3. Example report generated by N-Power AI for the Paired t-test scenario

**Power Analysis Report**

**1. Objective description:** The objective of this study is to investigate the effectiveness of a new medication in reducing cholesterol levels in patients aged 60 or above.

**2. Hypothesis:** The primary hypothesis of this study is that the new medication will reduce cholesterol levels after treatment compared to before treatment. Specifically, the null and alternative hypotheses are:

H0: μd = 0 (no reduction in cholesterol levels) H1: μd < 0 (reduction in cholesterol levels)

**3. Statistical Method/Analysis Plan:** The statistical method used for this study is a paired t-test, which compares the mean difference in cholesterol levels before and after treatment.

**4. Assumptions and Sample Size Calculations:** The assumptions made in this power analysis are:

- Effect size (d): -0.3947368
- Significance level: 0.05
- Power: 0.8
- Correlation coefficient between before and after measurements: 0.5

The calculated sample size required to achieve adequate power for the study is 41 pairs of measurements.

**5. Explanation of the power analysis results:** The power analysis results indicate that a sample size of at least 42 pairs of measurements (rounded up from 41.06547) would be necessary to detect a reduction in mean cholesterol levels after receiving the new medication with 80% power at a significance level of 0.05.

**6. Power Analysis Table:**

| Test type | Significance level One-Sided | Power | Sample size Pairs |
|---|---|---|---|
| Paired t-test | 0.05 | 0.8 | 42 |

Fig S4. Example report generated by N-Power AI for the One-way ANOVA scenario

**Power Analysis Report**

**1. Objective description:** The objective of this study is to investigate the effectiveness of four different teaching methods on improving standardized math scores in local classrooms.

**2. Hypothesis:** The primary hypothesis of this study is that there will be a significant difference in the mean score of students taught using different teaching methods, with the alternative hypothesis being that at least one group mean is different from the others.

**3. Statistical Method/Analysis Plan:** A balanced one-way analysis of variance (ANOVA) will be used to compare the mean scores of students across the four teaching method groups.

**4. Assumptions and Sample Size Calculations:** The assumptions made in this power analysis are that the sample means for each group will be 550, 598, 598, and 646, respectively, with a common standard deviation of 80. To achieve 80% power at a significance level of 0.05, the calculated sample size required for each group is 12.37865.

**5. Explanation of the Power Analysis Results:** The results indicate that to detect a difference in the mean score between groups with 80% power at a significance level of 0.05, a minimum of 13 students (rounded up from 12.37865) will be needed in each group.

**6. Power Analysis Table:**

| Test Type | Significance Level | Power | Groups | Sample Size |
|---|---|---|---|---|
| One-way ANOVA | 0.05 | 0.8 | 4 | 13 |

Fig S5. Example report generated by N-Power AI for the Chi-square test of independence scenario

**Power Analysis Report**

**Objective Description** The objective of this study is to assess gender differences in the severity of depression among college students.

**Hypothesis** The primary hypothesis of this study is that there is a significant association between depression severity and gender. The null hypothesis (H0) states that there is no association between depression severity and gender, while the alternative hypothesis (H1) states that there is an association between depression severity and gender.

**Statistical Method/Analysis Plan** The statistical method used in this study will be a chi-square test of independence to assess the association between depression severity and gender. The sample proportions for each depression severity group will be compared between females and males.

**Assumptions and Sample Size Calculations** The assumptions made in this power analysis are that the sample proportions for each depression severity group will be 0.6759, 0.1559, 0.1281, 0.0323, and 0.0078 in females, and will be 0.6771, 0.1519, 0.1368, 0.0241, and 0.0101 in males. The significance level is set at 0.05, and the desired power is 80%. Based on these assumptions, the calculated sample size required to achieve adequate power for the study is approximately 13069 students.

**Explanation of the Power Analysis Results** The results of the power analysis indicate that a minimum total number of 13069 students are needed to achieve 80% power for detecting the association between depression severity and gender at the significance level of 0.05. This means that if we collect data from approximately 13069 students, we will have an 80% chance of detecting a statistically significant association between depression severity and gender.

**Power Analysis Table**

| Test Type | Significance Level | Power | Sample Size |
|---|---|---|---|
| Chi-square test of independence | 0.05 | 0.8 | 13069 |

Reference

1. Daniel WW, Cross CL. Biostatistics: a foundation for analysis in the health sciences. Wiley; Nov 13, 2018.

2. UCLA: Statistical Consulting Group. Data Analysis Examples. Accessed Oct. 03, 2024. (https://stats.oarc.ucla.edu/other/dae/)

3. Gao W, Ping S, Liu X. Gender differences in depression, anxiety, and stress among college students: a longitudinal study from China. Journal of Affective Disorders. Feb 15, 2020;263:292-300.

4. Rosner BA. Fundamentals of biostatistics. Boston, MA: Brooks/Cole, Cengage Learning; 2015.
